# A conformationally constrained synthetic peptide efficiently inhibits malaria parasite entry into human red blood cells

**DOI:** 10.1101/2023.06.23.546305

**Authors:** Anamika Biswas, Akash Narayan, Suman Sinha, Kalyaneswar Mandal

## Abstract

Restricting the conformational freedom of a peptide by backbone cyclization and incorporation of an additional disulfide bond leads to a unique cyclic peptide that inhibits the invasion of red blood cells by malaria parasites efficiently. The engineered peptide exhibits twenty fold enhanced affinity towards its receptor (*Pf*AMA1) compared to the native peptide ligand (*Pf*RON2).

---

Protein-protein interactions (PPIs) play a crucial role in number of biological events, like cell-signalling, host pathogen interactions etc. ^1–4^ Targeting disease related PPIs with a small protein or peptide-based inhibitor, as opposed to a small molecule drug or a large protein biologic, has several advantages. ^5^ First, unlike small molecule drugs, small proteins or peptides provide a wide binding interface area, nearly 600-2000 Å^2, 4^ which resemblances the area of a PPI “hot-spot”. ^1,3,6^ As a result, a peptide or a small protein having a defined secondary or tertiary structure leads to high specificity and less off-target effect. When compared to a small molecule drug, accumulation of a peptide or small protein molecule in tissues is insignificant. ^4,7,8^ On the other hand, unlike a large protein biologic, like antibody, a small protein/peptide inhibitor does not encounter issues associated with tissue permeability.^9^ Most importantly, peptides or small proteins can be prepared chemically; hence, cost effective and amenable to unique chemical modifications which may result in increased binding affinity of an inhibitory peptide/protein. ^10^ Herein, we describe the design and chemical synthesis of a conformationally constrained peptide-based inhibitor to disrupt a key PPI responsible for the *Plasmodium falciparum* parasite entry into human red blood cells (RBC) that causes the life-threatening infectious disease, malaria.

Malaria is a mosquito-borne disease caused by *Plasmodium* species, the deadliest of which is *Plasmodium falciparum*. According to the latest WHO report, a total of 247 million malaria cases and 619 000 malaria deaths were documented in 2021. ^11^ The unavailability of a fully effective malaria vaccine along with the immerging resistance of the frontline medicine ‘artemisinin combination therapy’, have made this disease difficult to contain. ^12–15^ Hence, we are in desperate need to develop a potential alternative anti-malarial therapeutic.

While making an entry to the host cell, all apicomplexan parasites follow a unique invasion strategy involving moving junction formation between the host cell and the parasite. ^9,16,17^ The strong interactions between two malaria parasite proteins, namely, Apical Membrane Antigen 1 (AMA1) and Rhoptry Neck Protein 2 (RON2), form the moving junction. ^17–21^ By disrupting the interactions between AMA1 and RON2 with an exogenous peptide or a small protein, the parasite entry inside the human RBCs can be prevented very efficiently. ^22^ However, till date, only a limited number of peptide/ mini-protein based inhibitors have been exploited to stop the interactions between *P. falciparum* proteins *Pf*AMA1 and *Pf*RON2. ^17,22–27^ Several among those peptides are obtained using phage display ^28–32^ and several others are derived from the native 39-residue *Pf*RON2 ectodomain. ^18,20,26,33,34^ The 39-residue polypeptide constitutes a small extracellular peptidic loop of the *Pf*RON2 membrane protein that interacts with *Pf*AMA1 very tightly during the moving junction formation. ^35–37^ The interacting sequence of the ectodomain of the *Pf*RON2 protein (*Pf*RON2_2021-2059_) has turned out to be highly conserved across the *P. falciparum* strains and nearly 60% conserved across the *Plasmodium* species. ^38,39^ As a consequence, an inhibitor derived from the conserved *Pf*RON2_2021-2059_ sequence would abrogate the possibility of developing drug-resistance, which is otherwise a common issue for any drug development campaign. We set out to rationally introduce conformational constraints into the 39-residue polypeptide backbone to achieve a novel peptide inhibitor that would possibly mimic and stabilize the bound form of the *Pf*RON2 extracellular loop with *Pf*AMA1 (Figure 1A). The *Pf*RON2_2021-2059_ peptide has a short (four-residue) anti-parallel beta-sheet containing disulfide bonded loop and a short (seven-residue) alpha-helix at the N-terminus, in its bound form with the *Pf*AMA1. However, the isolated *Pf*RON2_2021-2059_ peptide in solution, when not bound to the hydrophobic groove of the *Pf*AMA1, is expected to be highly flexible with a disordered random coil structure, as evident from its CD spectra (Figure-S1). In order to give rigidity to the disordered random coil structure, we incorporated one additional disulfide bond by replacing Ala2031 and Ser2059 of *Pf*RON2_2021-2059_ with two Cys residues. The C*α* - C*α* distance between Ala2031 and Ser2059 is closure to the C*α* - C*α* distance between two cysteine residues in a disulfide bond. In addition, to restrict the flexibility of the two termini, we cyclized the molecule by connecting the N- and the C-termini with a variable-length poly-Gly linker. In order to identify an optimum length of the linker we synthesized five cyclic analogues, varying the number of Gly-residues from two to six (Figure 1B).

**Figure 1:**
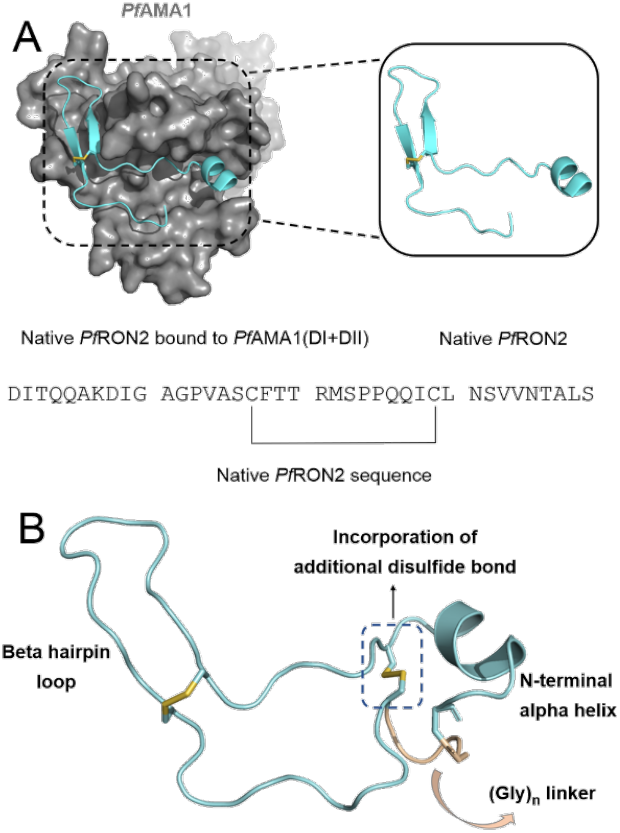
A. Native *Pf*RON2_2021-2059_ structure (cyan cartoon) in complexation with *Pf*AMA1(DI+DII) (gray surface); B. Designing cyclic nG2S inhibitors using *Pf*RON2_2021-2059_ as a template

All analogs of the *Pf*RON2_2021-2059_ peptide, except the acyclic one, were synthesized by Fmoc chemistry solid phase peptide synthesis (SPPS) as peptide hydrazide. The acyclic double-disulfide containing analog was made as carboxylic acid at the C-terminus. The cyclization of peptides containing C-terminal Leu-hydrazide and the free Cys_2059_ at the N-terminus were performed off-resin by thioesterification with MESNa followed by intra-molecular native chemical ligation (cyclization) in one-pot (Figure 2, S2). As it was an intra-molecular ligation, we found that aryl thiol catalyst was not needed, rather alkyl thiol MESNa was sufficient. Finally, all purified peptide analogs were allowed to fold to form disulfide bonds under air oxidation condition (Figure S2).

**Figure 2:**
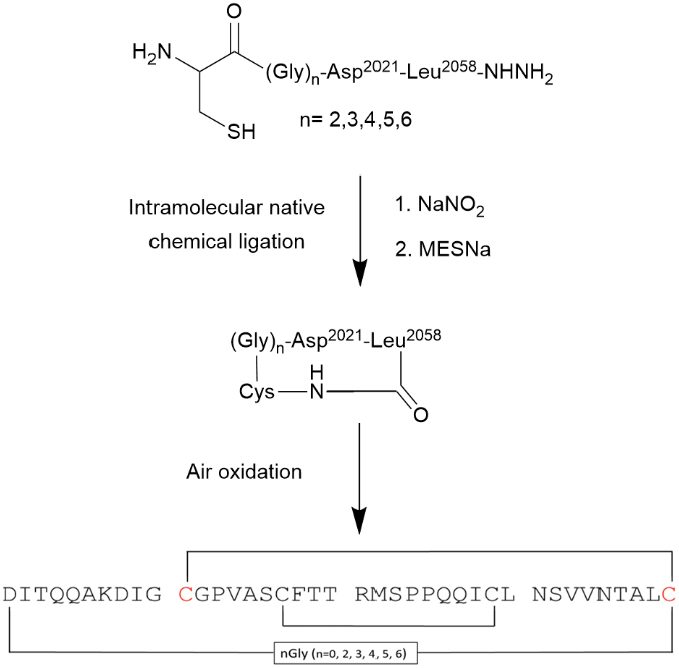
Synthetic scheme of cyclic nG2S using native chemical ligation

In order to identify the most potent candidate from the series of the designed *Pf*RON2_2021-2059_ mimetic inhibitors, we examined the functional activity by parasite growth inhibition assay (GIA) at a fixed concentration of 100 nM. Although, all the *Pf*RON2_2021-2059_ analogs stopped the merozoite invasion, the effect of the acyclic (RON2-2S) analog was minimum. This could be attributed to the fact that just the additional disulfide bond is not sufficient enough to restrict the flexibility of the two termini. Hence, the critical residues from the N-terminal helix ^20,40^ part are not always in the proximity of the *Pf*AMA1 D-II loop to make favourable contacts.

In contrary, all the backbone-cyclized analogues were found to enhance the binding activity to a significant extent. This observation further indicates that the cyclization probably reinforced the inhibitor to retain the native conformation of the *Pf*RON2_2021-2059_ peptide in its bound form with *Pf*AMA1, where the interacting residues from N-terminal helix can make enough contacts with the DII loop of the *Pf*AMA1 to further stabilise the protein-protein interactions.

Among all the cyclic peptides, 4G2S (Figure 3B) was found to be the most effective inhibitor with relatively high functional activity in GIA (Figure 3A). Hence, 4G2S peptide was chosen for further evaluation of its binding affinity against refolded *Pf*AMA1(DI+DII) ^41^ by SPR (Figure 4, Table S1). From the trend of GIA as well as SPR, it was confirmed that 4G2S has better binding affinity towards *Pf*AMA1 with respect to the native *Pf*RON2_2021-2059_. Hence, a full scan in GIA with a series of different concentrations of analytes were performed (Figure 3C, S3) to obtain its IC_50_ value. A leap of potency (> two-fold) of the 4G2S (IC_50_ ≈95 nM) analogue was observed when compared with the native *Pf*RON2_2021-2059_ (IC_50_ ≈220 nM) in disrupting the merozoite invasion into the RBC (Figure S3, Table 1). Moreover, when the association and dissociation rate of the most active 4G2S was compared with the native one (*Pf*RON2_2021-2059_), it was observed that the association rate of the 4G2S was similar to that of *Pf*RON2_2021-2059_, however, the dissociation rate was 20-fold lesser than the native peptide, which implies the increased residence time of the 4G2S analogue in complexation with the *Pf*AMA1(DI + DII) (Table 1).

**Table 1:** Comparison of functional activity of *Pf*RON2_2021-2059_ and its 4G2S analog measured by SPR (average of three repetitions) and GIA (average of two repetitions)

| Ligand | K<sub>D</sub><br>(nM) | K<sub>a</sub><br>(10<sup>5</sup> M<sup>-1</sup>.Sec<sup>-1</sup>) | K<sub>d</sub><br>(10<sup>-3</sup> Sec<sup>-1</sup>) | % Inhibition at<br>100nM in GIA | IC<sub>50</sub> Value<br>(nM) |
| --- | --- | --- | --- | --- | --- |
| *Pf*RON2<sub>2021–2059</sub> | 21.60 ± 0.64 | 3.61 ± 0.77 | 7.83 ± 1.88 | 11.66 | 220 |
| 4G2S | 1.11 ± 0.23 | 3.40 ± 0.70 | 0.37 ± 0.04 | 61.71 | 95 |

**Figure 3:**
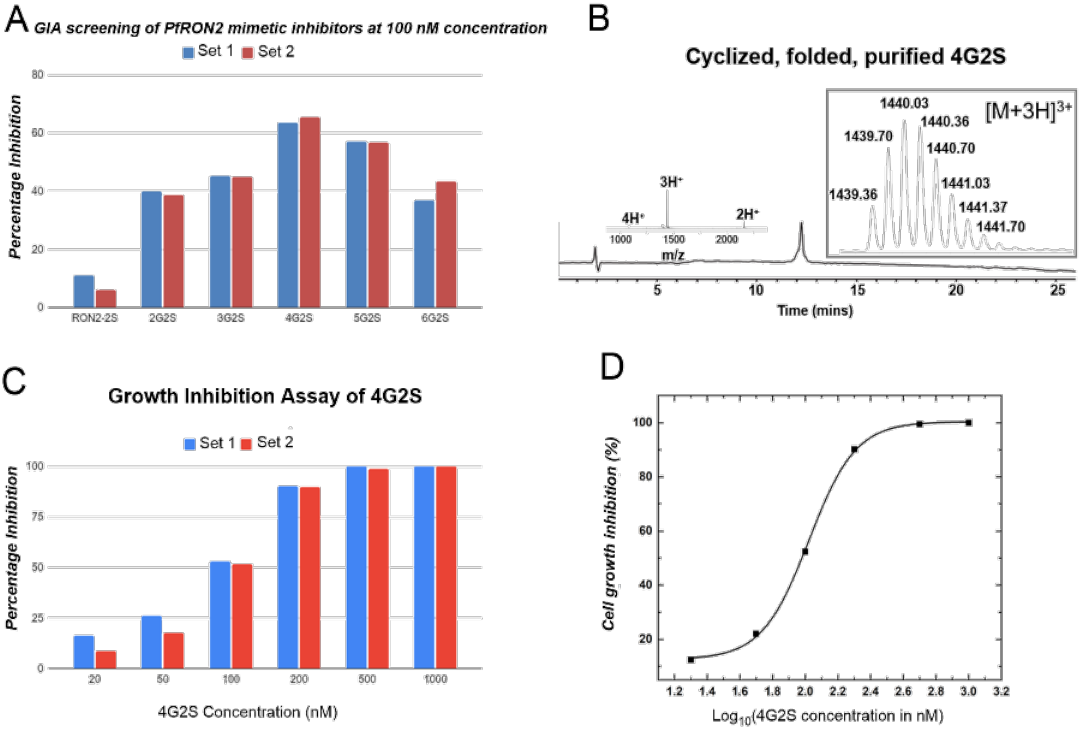
A. Functional activity of all peptide analogs at 100 nM concentration by GIA screening (performed twice for data reproducibility); B. LC-MS chromatogram of purified folded cyclic 4G2S (observed mass (most abundant isotopologue) = 4317.07 ± 0.02 Da, calculated mass: 4317.00 Da); C. Full scan of 4G2S in GIA to check its percentage inhibition in different concentrations (performed twice for data reproducibility); D. Cell growth inhibition profile of 4G2S in log scale to obtain the IC_50_ value (average of two sets).

**Figure 4:**
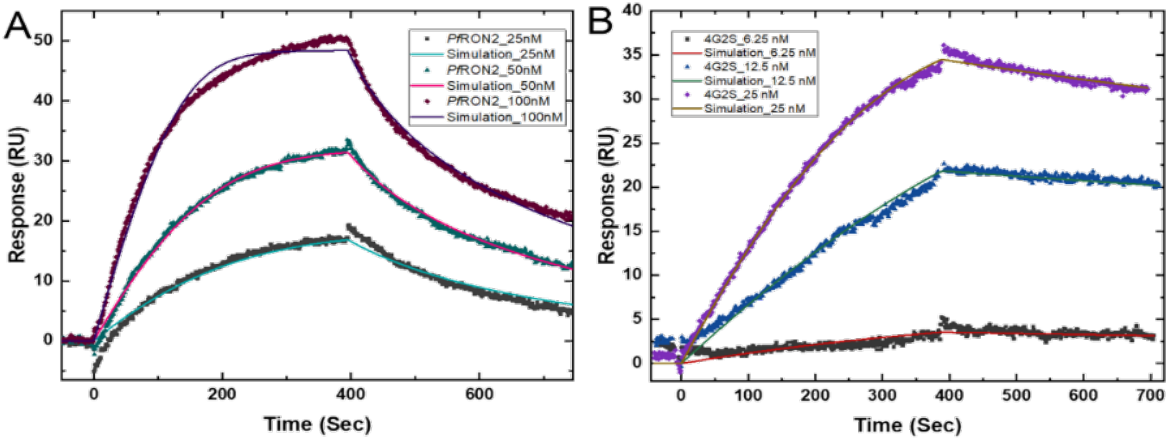
Representative SPR sensorgram (one of three repeats) of A. *Pf*RON22021-2059 and B. 4G2S, to compare the binding affinity of the analytes towards *Pf*AMA1(DI+DII).

Moreover, when the functional activity of 4G2S was further checked by GIA, it was observed that 4G2S successfully stopped the merozoite invasion to the human RBC with an IC_50_ value of ∼95 nM (Figure 3C, D), whereas *Pf*RON2_2021-2059_ does the same with an IC_50_ value of ∼220 nM. Therefore, both the SPR and GIA experimental data firmly indicated that 4G2S is a two-fold more potent inhibitor with 20-fold higher binding affinity than *Pf*RON2_2021-2059_. In order to investigate the apparent mode of binding of the 4G2S analog, we performed molecular docking using ZDock software. From the molecular docking study, we found that the energy minimized docked structure closely resembled the *Pf*AMA1 bound form of the *Pf*RON2_2021-2059_ peptide with slight deviation at the helix region (Figure 5).

**Figure 5:**
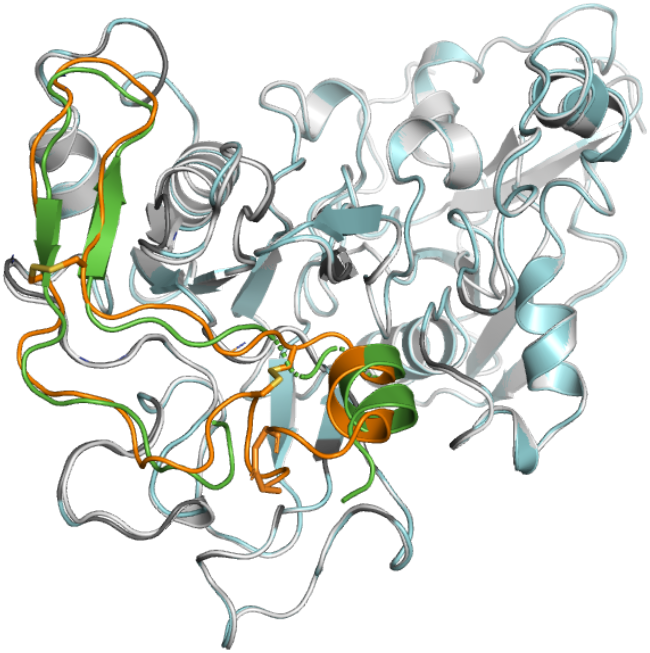
Superimposition of the energy minimized molecular docking structure of 4G2S with *Pf*RON2_2021-2059_ in complexation with *Pf*AMA1(DI+DII). (*Pf*RON2_2021-2059_ (green) in complexation with *Pf*AMA1 (gray); 4G2S (orange))

In summary, a potential cyclic peptide inhibitor has been designed based on sequence of the well-conserved *Pf*RON2_2021-2059_ peptide. In our study, 4G2S, successfully stopped the merozoite invasion of the human RBC with an IC_50_ value of ∼95 nM and it showed a very high affinity towards the *Pf*AMA1(DI+DII) (K_D_ ∼1.1 nM). To the best of our knowledge,, 4G2S is the most potent inhibitor based on the 39 mer native *Pf*RON2_2021-2059_ sequence reported so far. Most importantly, the superimposed docked structure of *Pf*RON2_2021-2059_ and 4G2S in complexation with *Pf*AMA1(DI+DII) (Figure 5) also implied that by doing the cyclization and incorporation of an additional disulfide bond to the backbone we could possibly able to mimic the conformation of the *Pf*RON2 ectodomain, leading to the enhanced residence time of 4G2S when in complex with the *Pf*AMA1.

## Acknowledgements

This work was supported by the DBT/Wellcome Trust India Alliance Grant (grant no. IA/I/15/1/501847) awarded to K.M. and the intramural funds at TIFR Hyderabad from the Department of Atomic Energy (DAE), Government of India, under Project Identification No. RTI-4007.

