## Supplementary file for "A conformationally constrained synthetic peptide efficiently inhibits malaria parasite entry into human red blood cells"

### General Methods

#### 1. Materials and methods

All chemicals, used in this project, were obtained from commercial sources and used without further purification. All peptides were synthesized using an automated peptide synthesizer (Tribute-UV/IR from Protein Technologies, USA). Analytical reverse-phase HPLC was performed on an Agilent HPLC instrument using an Agilent zorbax SB-C3 (5 µm), 4.6×150 mm reverse-phase silica column at a flow rate of 0.9 mL/min using 0.1% TFA in $H_2O$ as solvent A and 0.08% TFA in acetonitrile as solvent B. The UV absorbance of the column eluent was monitored at 214 nm wavelength. The peptide masses were measured across the peak by on-line LC-MS using an Agilent 1290 infinity II/6530 Q-TOF LC/MS instrument. The deconvolution of the charge states of the observed mass was carried out using Agilent MassHunter Qualitative Analysis software (version B.07.00), and the deconvoluted mass of the most abundant isotopologue has been reported. Preparative reverse-phase HPLC (RP-HPLC) of crude peptides was performed with a Waters 1525 preparative HPLC system using Waters C4 (5 µm, 300 Å, 10 x 250 mm) or Agilent ZORBAX-SB C3 (5 µm, 80 Å, 9.4 x 250 mm) columns at 40 °C using an appropriate shallow gradient at a flow rate of 5 mL/min using solvent A and B. Fractions containing the purified target peptide were identified by ESI-MS. Selected pure fractions were then pooled and lyophilized.

#### 2. Circular dichroism spectra of *Pf*RON2$_{2021\text{-}2059}$

The CD spectra of chemically synthesized *Pf*RON2$_{2021\text{-}2059}$ was recorded using a Jasco CD spectrometer. For measuring the spectrum over the wavelength range of 185-260 nm, sample (~0.4 mg/ml, based on the dry weight) was dissolved in water and water - Trifluoroethanol

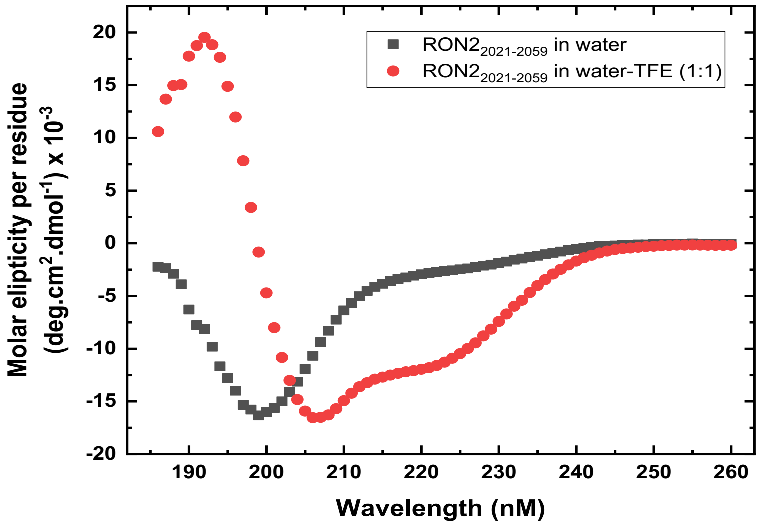

***Figure S1: CD spectra of PfRON2$_{2021\text{-}2059}$ with and without TFE***

(1:1) separately, and transferred to a Hellma cuvette of pathlength 1 mm. The spectrum was recorded at 25°C. Molar ellipticity per residue, was calculated using the following equation,

Molar ellipticity (deg.cm$^2$.dmol$^{-1}$) = $\frac{Elipticity\ (mdeg)*Mw}{10*c*l*Nr}$

Where, Mw is the molecular weight of the protein, c is concentration in g/l, l is the pathlength of the cuvette in cm and Nr is the number of amino acid residues in the protein.

#### 3. Expression, purification, and refolding of *Pf*AMA1(DI+DII)

Refolded *Pf*AMA1(DI+DII) was obtained by recombinant expression of the protein in *E. coli* BL21 (DE3) RIL cell using the previously published protocol. [1] The expression plasmid encoding 3D7 *Pf*AMA1(DI+DII) (104-438) sequence was obtained from GenScript (NI, USA). The protein contains an N-terminal hexa-histidine tag which is required for purification of the protein by Ni-affinity chromatography. The same His-tag was also used for immobilizing the refolded protein in Ni-NTA functionalised chip for SPR experiments.

#### 4. Synthesis, cyclisation, and oxidative folding of *Pf*RON2$_{2021\text{-}2059}$ and other *Pf*RON2$_{2021\text{-}2059}$ mimetic peptide analogs

Using an automated peptide synthesizer, the 39-mer *Pf*RON2$_{2021\text{-}2059}$ (Asp$_{2021}$IleThrGlnGlnAlaLysAspIleGlyAlaGlyProValAlaSerCysPheThrThrArgMetSerProProGln-GlnIleCysLeuAsnSerValValAsnThrAlaLeuSer$_{2059}$) was synthesized at an elevated temperature

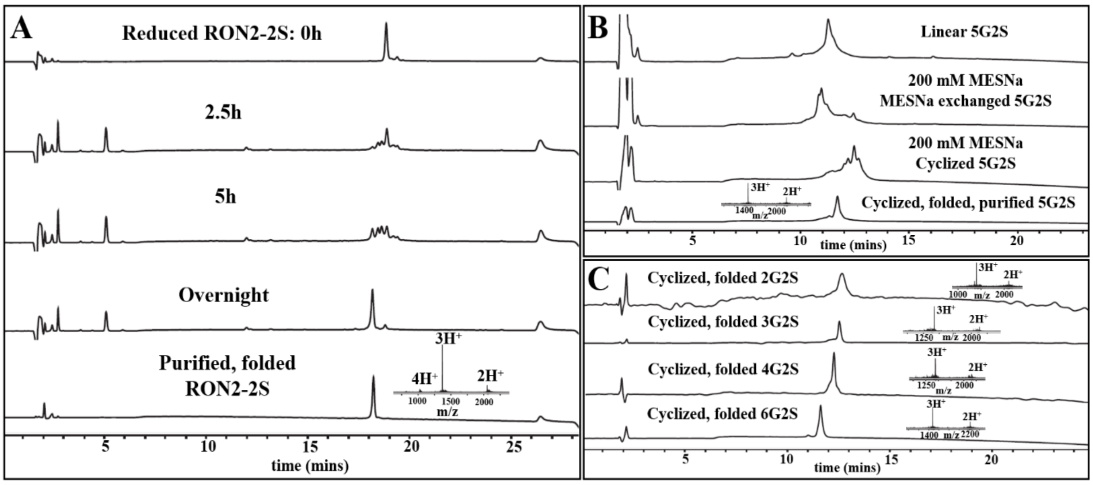

***Figure S2: A. Folding profile of acyclic PfRON2-2S by air oxidation condition (observed mass = 4106.91 ± 0.03 Da; calculated mass: 4106.92 Da); B. HPLC chromatogram showing one-pot cyclisation of crude 5G2S followed by folding. The ESI-MS data is shown inset (observed mass (most abundant isotopologue) = 4374.01 ± 0.03 Da; calculated mass: 4374.02 Da); C. LC-MS chromatogram of other cyclized, folded nG2S peptide analogs [2G2S (observed mass (most abundant isotopologue) = 4202.97 ± 0.03 Da; calculated mass: 4202.96 Da), 3G2S (observed mass (most abundant isotopologue) = 4260.02 ± 0.02 Da; calculated mass: 4259.98 Da), 4G2S (observed mass (most abundant isotopologue) = 4317.07 ± 0.02 Da, calculated mass: 4317.00 Da), 6G2S (observed mass (most abundant isotopologue) = 4431.03 ± 0.03 Da; calculated mass: 4431.04 Da)].***

in a single stretch on 2-cholorotrityl chloride resin using Fmoc protected amino acids by step-wise solid phase peptide synthesis following the reported protocol. [2] The other *Pf*RON2-analogs, *Pf*RON2-2S (Asp$_{2021}$IleThrGlnGlnAlaLysAspIleGlyCysGlyProValAlaSerCysPheThrThrArgMetSerProProGlnGlnIleCysLeuAsnSerValValAsnThrAlaLeuCys$_{2059}$) and nG2S (Cys$_{2059}$[Gly]$_n$Asp$_{2021}$IleThrGlnGlnAlaLysAspIleGlyCysGlyProValAlaSerCysPheThrThrArgMetSerProProGlnGlnIleCys-LeuAsnSerValValAsnThrAlaLeuNHNH$_2$), where n = 2-6, were also synthesized following the same protocol on a 2-cholorotrityl chloride resin. After the synthesis, peptides were cleaved from the solid support followed by reverse-phase HPLC purification. The peptide hydrazides (for the cyclic inhibitors) were dissolved in 6M Gu.HCl, 0.2M phosphate buffer (pH ~3.5) and further oxidized with 40 mM sodium nitrite. We then cyclized nG2S analogs in presence of 200 mM MPAA/MESNa at pH ~6.8. After the completion of the cyclization, monitored by LCMS, cyclized peptides were purified by reverse-phase HPLC and lyophilized. Lyophilized peptides (*Pf*RON2$_{2021\text{-}2059}$ and all its analogs) were folded under air oxidation condition (at pH 8.4, same as *Pf*RON2$_{2021\text{-}2059}$, [1]) followed by purification and lyophilization. Lyophilized folded peptides were stored at 4°C and further used to study their functional activity by SPR and parasite growth inhibition assay (GIA).

#### 5. GIA of chemically synthesized *Pf*RON2$_{2021\text{-}2059}$ and its analogous peptides

For the GIA experiment, tight synchronization of *Plasmodium falciparum* 3D7 cell line culture was carried out beforehand. Synchronized intraerythrocytic schizont stage parasites with 0.08% parasitemia were allowed to grow in nutrient rich media after treating with a particular concentration (100 nM) of chemically synthesized peptides (RON2-2S, 2G2S, 3G2S, 4G2S, 5G2S and 6G2S) for two cycles (55h) in a parallel set up following the previously described procedure. [3] Once the incubation period ends, 2 µL pallet culture was taken after centrifugation to prepare thin smear for performing standard Giemsa counting assay. The slides containing thin smears of each of the treated and control culture were stained with Giemsa after fixation with methanol. The slides were observed under microscope using 100X oil objective with 0.5X camera adapter and the images were taken from the random adjacent fields for the further processing. Around 4000 RBCs including the infected RBCs (iRBCs) were

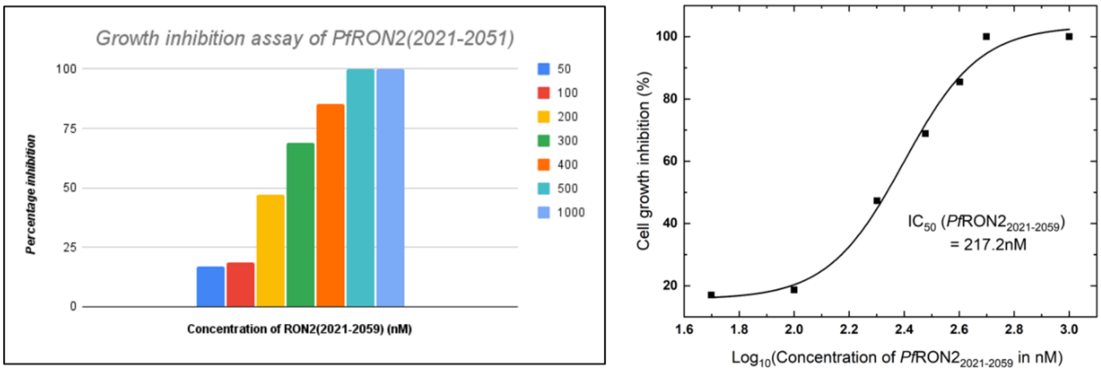

***Figure S3: Growth inhibition assay of PfRON2$_{2021\text{-}2059}$ to obtain the IC$_{50}$ value***

counted using auto counting software (followed by cross-checking manually) and final parasitemia was calculated. This assay was performed twice for the data reproducibility.

To obtain the $IC_{50}$ value, GIA experiment was performed for 4G2S in different concentrations (20, 50, 100, 200, 500, 1000 nM) and compared with *Pf*RON2$_{2021-2059}$ [1,4] (Figure S3, 3C).

#### 6. Comparing the binding affinity of the peptides against *Pf*AMA1(DI+DII) by using surface plasmon resonance

The binding affinity of the *Pf*RON2$_{2021-2059}$ analogous peptides were measured and compared with *Pf*RON2$_{2021-2059}$ against *Pf*AMA1(DI+DII) using BI-4500AP SPR instrument. 2.4 µM freshly refolded N-terminal His-tagged *Pf*AMA1(DI+DII) was immobilized at pH 7.4 in a Ni-NTA functionalized chip using 10mM PBS as running buffer with 0.005% tween-20. Out of the four parallel flow cell channels, three were immobilized with *Pf*AMA1 protein for data reproducibility while the other one was kept free as reference channel by injecting only the running buffer. At this point, to avoid non-specific interactions, 40 µM EDTA was added to the running buffer. Lyophilized peptide analytes were dissolved in water and diluted serially with the running buffer to prepare a series of analyte concentrations (6.25 nM, 12.5 nM, 25 nM). The analytes were passed through the flow cell at 30 µl/min flow rate and allowed to interact with the immobilized *Pf*AMA1 protein for 360 sec to get the association curve. Once the saturation point is reached, only running buffer was passed through the flow cell in the same flow rate to measure dissociation kinetics. To obtain the final sensorgram, reference channel was subtracted and the binding profiles were fitted using 1:1 Langmuir adsorption binding isotherm with the help of BI-data analysis software.

***Table S1: Summery of the binding experiments performed by SPR in multiple repetitions***

| Analyte | Repetitions | $K_D$ (nM) | $K_a$ ($10^5$ $M^{-1}.Sec^{-1}$) | $K_d$ ($10^{-3}$ $Sec^{-1}$) |
|---|---|---|---|---|
| *Pf*RON2 | Set 1 | 21.03 | 3.54 | 7.44 |
| | Set 2 | 21.29 | 2.70 | 5.75 |
| | Set 3 | 22.49 | 4.59 | 10.31 |
| 4G2S | Set 1 | 1.17 | 3.56 | 0.42 |
| | Set 2 | 0.81 | 4.16 | 0.34 |
| | Set 3 | 1.36 | 2.48 | 0.34 |

#### 7. Molecular Docking of cyclic 4G2S and *Pf*RON2$_{2021-2059}$ to understand the binding pose while in complexation with *Pf*AMA1(DI+DII)

Studies were conducted using ZDock [5] webserver. ZDOCK 3.0.2 [6,7] employs the Fast Fourier Transform algorithm to facilitate a thorough exploration of docking possibilities across a 3D grid. It incorporates a blend of shape complementarity, electrostatics, and statistical potential factors to evaluate and rank the results. Topmost pose from the docking calculations was subjected to energy minimization using Gromacs [8,9] software package. The steepest descent algorithm [10] was applied to optimize the system to its lowest possible energy.
